# A-to-I mRNA editing in a ferric siderophore receptor improves competition for iron in *Xanthomonas oryzae*

**DOI:** 10.1101/2020.12.03.409276

**Authors:** Wenhan Nie, Sai Wang, Jin Huang, Qin Xu, Peihong Wang, Yan Wu, Ayizekeranmu Yiming, Iftikhar Ahmad, Bo Zhu, Gongyou Chen

## Abstract

Adenosine-to-inosine (A-to-I) RNA editing, which is catalyzed by the adenosine deaminase RNA-specific family of enzymes, is a frequent post-transcriptional modification in metazoans. Research on A-to-I editing in bacteria is limited, and the importance is underestimated. In this study, we show that bacteria may use A-to-I editing as an alternative strategy to promote uptake of metabolic iron. The T408A editing event of *xfeA* in *Xanthomonas oryzae* pv. *oryzicola* (*Xoc*) senses extracytoplasmic iron and changes the hydrogen bonding network of ligand channel domains. The frequency of A-to-I RNA editing during iron-deficient conditions increased by 76.87%, which facilitated the passage of iron through the XfeA outer membrane channel. When bacteria were subjected to high iron concentrations, the percentage of A-to-I editing in *xfeA* decreased, which reduced iron transport via XfeA. Furthermore, A-to-I RNA editing increased expression of multiple genes in the chemotaxis pathway, including methyl-accepting chemotaxis proteins (MCPs) that sense concentrations of exogenous ferric enterobactin (Fe-Ent) at the cytoplasmic membrane. A-to-I RNA editing helps *Xoc* move towards an iron-rich environment and supports our contention that editing in *xfeA* facilitates entry of a ferric siderophore. Overall, our results reveal a new signaling mechanism that bacteria use to facilitate iron uptake and improve their competitiveness.

## INTRODUCTION

Iron is one of the most abundant elements in the Earth’s crust and is essential for all life forms due to its roles in respiration, photosynthesis, DNA replication, oxygen transport and protection from various stresses (Skaar, 2010; L. Wang et al., 2016). Bacterial pathogens acquire iron within the host to promote survival and replication. The ability to successfully compete for iron is critical for bacterial pathogens that invade hosts when iron is limiting, which is a form of nutritional immunity (Hood & Skaar, 2012). Animal and plant hosts can retain iron by sequestering the element with various proteins or low molecular weight compounds (Fischbach, Lin, Liu, & Walsh, 2006; Kehl-Fie & Skaar, 2010), whereas bacteria have developed efficient iron uptake mechanisms based on high-affinity siderophores (Hider & Kong, 2010; Raines et al., 2016).

Many gram-negative bacteria such as *Xanthomonas* utilize siderophores to scavenge iron (Ryan et al., 2011). Siderophores are generally synthesized within the bacterial cell and then secreted to the extracellular milieu where they capture Fe^3+^. The resulting ferric siderophore complex is recognized at the outer membrane by TonB-dependent receptors (TBDR) and then actively transported into the periplasm of gram-negative species (Schalk, Mislin, & Brillet, 2012). One of the most well-characterized TBDRs is FepA, which transports the siderophore enterobactin (Ent) into the *Escherichia coli* periplasm. In the periplasm, ferrienterobactin (Fe-Ent) is sequestered by the periplasmic binding protein FepB, which transfers Fe-Ent to the inner membrane for further transport into the cytoplasm via FepCD (Raines et al., 2016). In the cytoplasm, Fe-Ent is hydrolyzed to release Fe^3+^ and converted to Fe^2+^ for further usage in the cell (Raymond, Dertz, & Kim, 2003).

RNA editing involves the alteration of ribonucleic acid after the molecule is produced by RNA polymerase and may involve deletion, insertion or base substitution events. One of the more common RNA editing events is the deamination of adenosine to inosine (A-to-I), which is catalyzed by the dsRNA-specific adenosine deaminase (ADAR) family of enzymes (Yablonovitch, Deng, Jacobson, & Li, 2017). The conversion of adenosine to inosine destabilizes dsRNA base pairing, interferes with the RNAi pathway and can change the amino acid sequence of the resulting protein. Post-transcriptional A-to-I editing has proven important in eukaryotes where it can drive adaptive evolution of the host (Yablonovitch et al., 2017); however, little is known about the incidence and function of A-to-I RNA editing in bacteria (Bar-Yaacov et al., 2017).

Our lab is interested in the role of A-to-I RNA editing in *Xanthomonas oryzae* pv. *oryzicola* (*Xoc*), which causes bacterial leaf streak in rice. We previously identified an A-to-I mRNA editing event in *Xoc* designated T408A (Nie et al., 2020); this editing event changed threonine to alanine in residue 408 of a ferric siderophore outer membrane receptor (FepA orthologue). In this study, the role of the T408A editing event was explored in *xfeA*, the *Xoc* homolog of *fepA*. The results show that the T408A editing event in *xfeA* enhances bacterial iron uptake capacity and improves tolerance to iron-limiting conditions.

## RESULTS

### T408A editing in *xfeA* is dependent on iron concentrations

Editing in *xfeA* was analyzed in cDNA samples of *Xoc* BLS256 grown in media amended with the iron chelating agent 2,2’-dipyridyl (DP) or supplemented with FeCl_3_ (Fig. 1a). The results suggested that T408A editing was dependent on available iron, and the incidence of editing was higher as the concentration of DP increased and iron became limiting (Fig. 1a). When *Xoc* BLS256 was cultured in non-amended NB, only 21.27% of the cDNAs contained the A-to-I editing event (A→G, Fig. 1b). Similarly, very low levels of editing (1.06%) were observed in BLS256 grown in NB plus 100 μM FeCl_3_ (Fig. 1a). In samples subjected to iron chelation, 37.39%, 76.87% and 78.03% of the cDNA samples exhibited A→G editing in the presence of 50, 100 and 150 μM DP, respectively (Fig. 1a,b). To block RNA editing, the synonymous mutation (ACG → ACA) was generated in the *Xoc* mutant T408^silent^, and no editing was detected in *xfeA* (Fig. 1b, red arrow).

**Figure 1.**
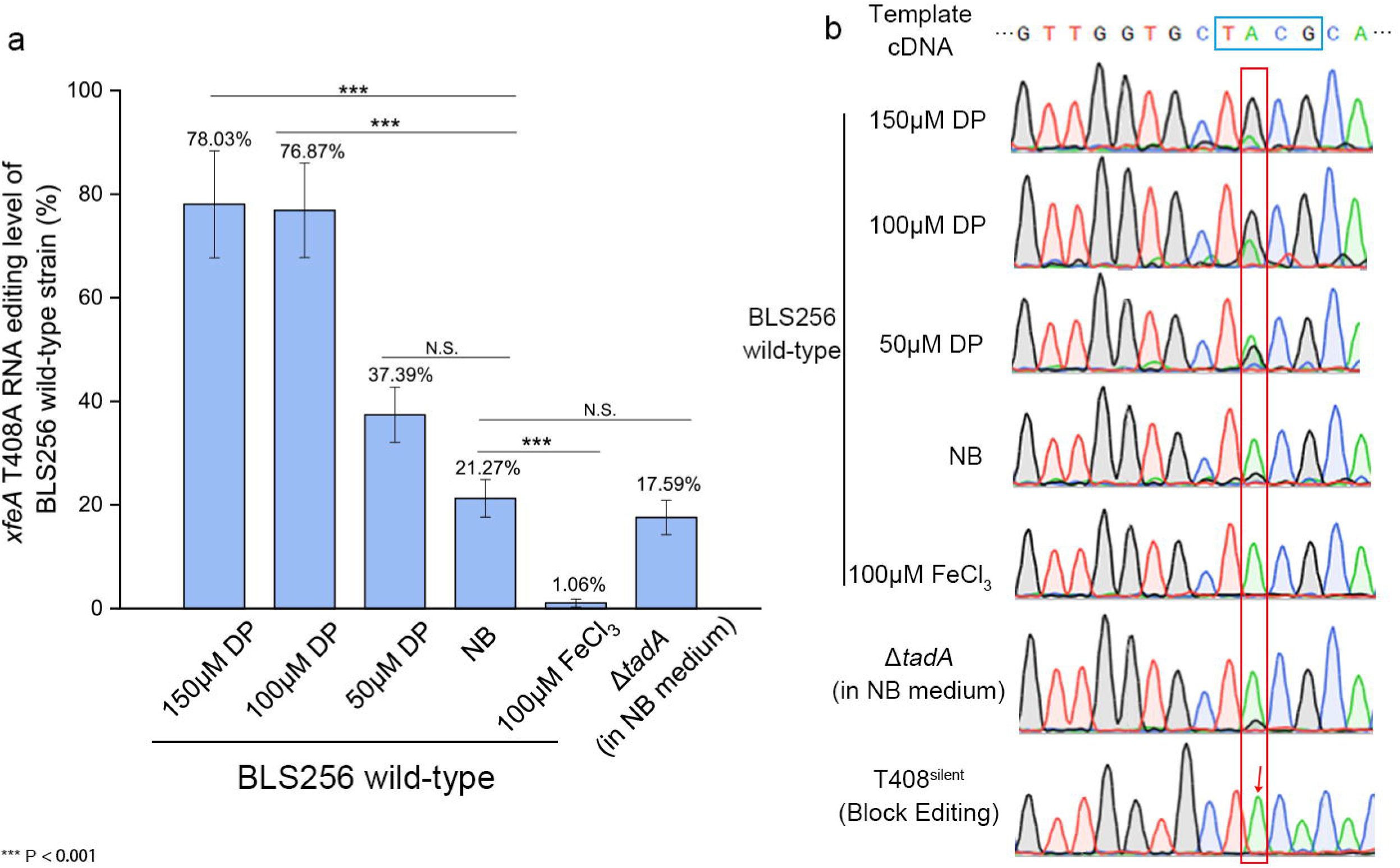
Analysis of A-to-I RNA editing in *xfeA* (T408A). (a) Percentage of editing as determined by sequence analysis of cDNA in *Xoc* BL256 (wild-type, WT) grown in NB, NB+50 μM 2,2’-dipyridyl (DP, iron chelator), NB+100 μM DP, NB+150 μM DP, and NB+100 μM FeCl_3_. The *Xoc ΔtadA* mutant containing a deletion in the gene encoding tRNA-specific adenosine deaminase is also included and was cultivated in NB. PCR fragments were obtained from cDNA derived from mRNA using primers described in Methods. Samples were incubated overnight to log phase and then subjected to Sanger sequencing. Three independent biological replicates were carried out in this study. ***, significant at *P* < 0.001; N.S., no significance. (b) Chromatograms reveal editing in different *Xoc* strains and culture conditions. The edited region, TACG, is framed by the blue rectangle in the template cDNA, and the percentage of bases edited from A (green) to G (black) is represented by peak height inside the red rectangle. In the chromatogram of the T408^silent^ strain, the red arrow indicates where A-to-I editing is blocked.

Editing was also compared by RNA-seq analysis of the T408A (contains A→G point mutation) and T408^silent^ strains grown in NB medium and analyzed with AIMAP (A-to-I modification analysis pipeline) (S. Wang et al., 2020). The results indicated that *xfeA* editing was 100% and 0% in the T408A and T408^silent^ mutants, respectively (Table S4). These results confirmed that A-to-I RNA editing in *xfeA* was blocked in the T408^silent^ mutant and emphasized the importance of the editing motif. A-to-I editing in *xfeA* occurred at low levels in the Δ*tadA* mutant (Fig. 1a), indicating that editing was not dependent on *tadA*-encoded adenosine deaminase activity; this was surprising since the TACG motif and the RNA secondary structure of *xfeA* contains motifs and structures recognized by TadA (Bar-Yaacov et al., 2017) (Fig. 1b, S2).

### T408A mutants show increased tolerance to iron depletion

Growth of the *Xoc* BL256 (WT), T408A, and T408^silent^ strains were compared in 0, 50, 100 and 150 μM DP (Fig. 2a-d). Strains grown in NB medium with or without 50 μM DP showed similar growth patterns (Fig. 2a and 2b); however, a longer lag phase and reduced growth rate were observed in WT and T408^silent^ strains grown in NB supplemented with 100 or 150 μM DP (Fig. 2c and 2d). The T408A strain (fixed, 100% editing) showed enhanced tolerance to DP and better growth than the WT and T408^silent^; the latter strain was severely impaired in growth at 150 μM DP due to lack of editing. The results indicate that T408A editing helps *Xoc* adapt to iron-deficient conditions.

**Figure 2.**
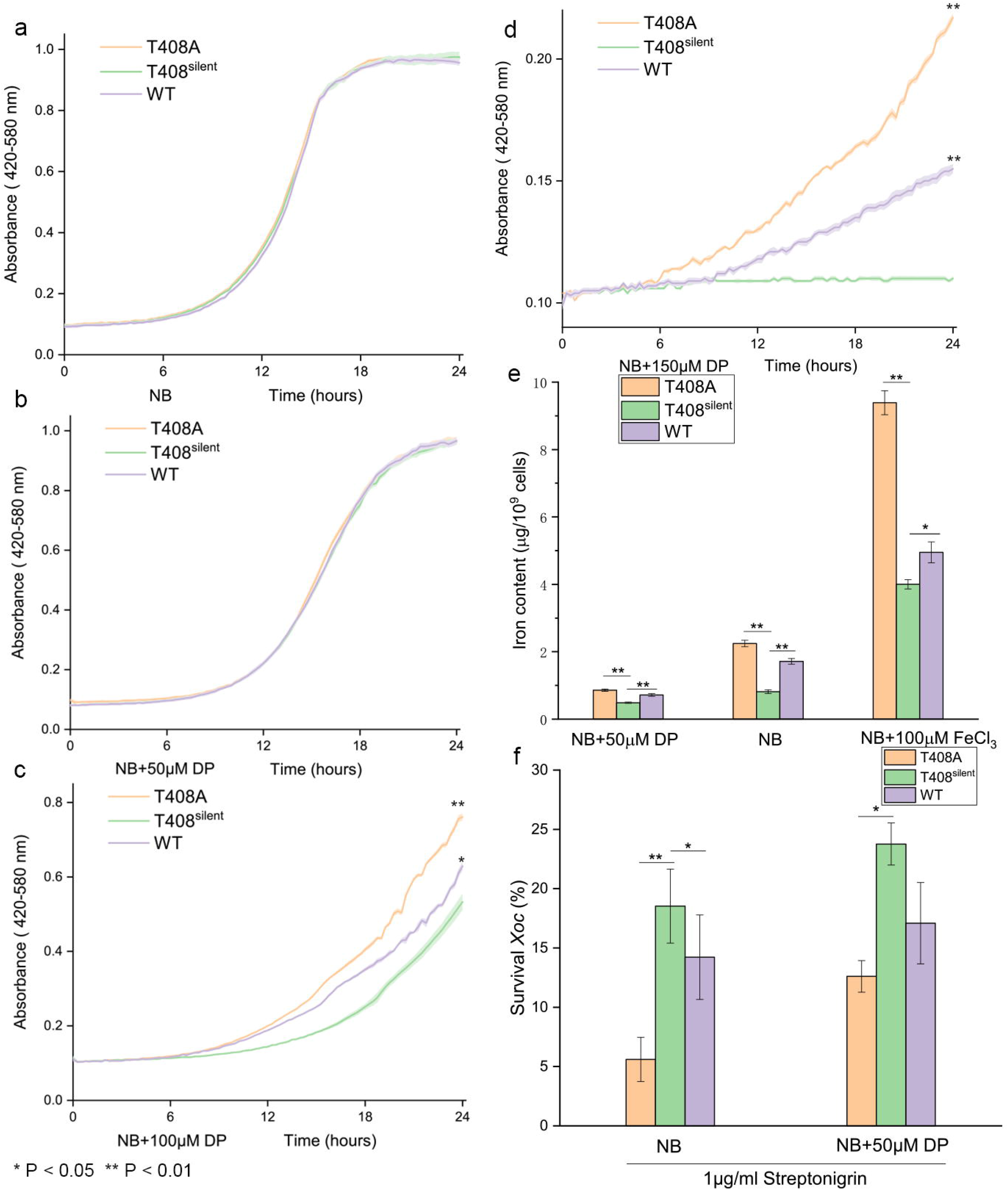
T408A RNA editing modulates bacterial responses to iron concentrations. (a) Growth curve of *Xoc* T408A, T408^silent^ and WT in NB medium. Strains were grown in quadruplicate to mid-exponential phase, diluted to OD_600_=0.1, transferred to fresh NB and monitored for growth in a Bioscreen C apparatus at 28°C. Growth in NB supplemented with 50 μM, (c) 100 μM, and (d) 150 μM DP. Error intervals (shaded regions bordering each line) indicate mean ± SE of four replicates. The F-test was used to compare growth of the T408A and WT strains with T408^silent^; * and ** indicate significance at *P* < 0.05 and 0.01, respectively. (e) Iron content in *Xoc* T408A, T408^silent^ and WT strains. Bacterial cells were cultured in NB, NB + 50 μM DP or NB + 100 μM FeCl_3_. The iron content in *Xoc* was measured with inductively-coupled plasma spectroscopy (ICP-OES). (f) Survival rates of *Xoc* T408A, T408^silent^ and WT strains exposed to streptonigrin. Strains were grown to mid-log phase (OD_600_=0.5) in NB or NB+DP and treated with 1 μg/ml streptonigrin for 24 h. Survival was assessed by colony counts from 10-fold serial dilutions. Error bars in (e) and (f) represent standard deviations (*n* = 3). * and ** indicate significant differences between the T408A or WT strains with T408^silent^ (control) at *P*<0.05 and 0.01, respectively (Student’s *t*-test).

### T408A editing enhances iron uptake

Intracellular iron concentrations were measured in the T408A, T408^silent^ and WT strains by inductively coupled plasma optical emission spectrometry (ICP-OES). In iron-deficient conditions (NB medium + 50 μM DP), iron accumulation in *Xoc* T408A and WT cells strains was significantly higher than that in cells of the T408^silent^ strain (Fig. 2e). In non-amended NB, iron concentrations in the T408A and WT strains increased by 175.3% and 110.1%, respectively, as compared to the T408^silent^ strain. In iron-replete conditions (NB + 100 μM FeCl_3_), the iron content of T408A and WT cells was 145.6 and 23.6% higher than T408^silent^, respectively (Fig. 2e). These results suggest that the T408A strain has better iron uptake than the WT and T408^silent^ strains in the presence of supplemental iron, which further confirms the importance of T408A RNA editing in *xfeA*.

Streptonigrin (SNG) is an antibiotic that requires iron for antibacterial activity, and its toxicity is correlated with intracellular iron concentrations (Yeowell & White, 1982). To further characterize the relationship between T408A editing and iron uptake, *Xoc* resistance to SNG was measured. When *Xoc* strains were exposed to 1 μg/mL SNG, survival of T408A and the WT were 69.8 and 23.2% lower than T408^silent^ strain, respectively. When the three strains were grown in iron-deficient conditions (NB + 50 μM DP), survival in response to SNG was improved and could be ranked as follows: T408A < WT < T408^silent^ (Fig. 2f). Collectively, these results supported our contention that *xfeA* T408A editing event facilitates iron uptake and increases tolerance to iron deficient conditions.

### T408A editing upregulates genes related to chemotaxis

RNA-seq was used to analyze expression in the T408A and T408^silent^ strains to further investigate the effect of *xfeA* T408A editing. The correlation coefficients (*r*) in two replicate experiments were 0.991 and 0.998, suggesting satisfactory reproducibility of RNA-seq data under the experimental conditions. Based on the cutoff values described in Methods, 138 differentially expressed genes (DEGs) were identified (Fig. 3a, Table S3). KEGG analysis indicated that the chemotaxis pathway was enriched in these DEGs (Table. S3). Interestingly, 21 of 24 chemotaxis pathway genes were up-regulated at least 1.5-fold [false discovery rate (FDR) < 0.01]. The following five genes were selected to verify the RNA-seq data by qRT-PCR: *xoc_2278* (encodes a two-component response regulatory protein similar to CheB); *xoc_2280* (encodes a chemotaxis methyltransferase similar to CheR); and *xoc_2289*, *xoc_2291*, *xoc_2297*, which are genes encoding methyl-accepting chemotaxis proteins (MCPs). *XfeA* was also included for comparative purposes. The results showed that expression levels of the five chemotaxis genes were correlated with iron concentration; in other words, expression was highest when NB was supplemented with 100 μM FeCl_3_ and lowest when the iron chelator DP was added to media (Fig. 3b). Expression of *xfeA* was relatively constant and was not impacted by iron levels (Fig. 3b; Fig. S1), and transcription of the chemotaxis genes was generally higher in T408A as compared to T408^silent^ (Fig. 3b). A diagram of the chemotaxis pathway was modeled using KEGG map entry 02030 (https://www.genome.jp/entry/map02030) and used to illustrate the upregulated genes due to T408A editing. Interestingly, almost all genes in the chemotaxis pathway were upregulated (Fig. 3c), which suggested that the T408A editing event may induce sensitivity to one or more chemoattractants.

**Figure 3.**
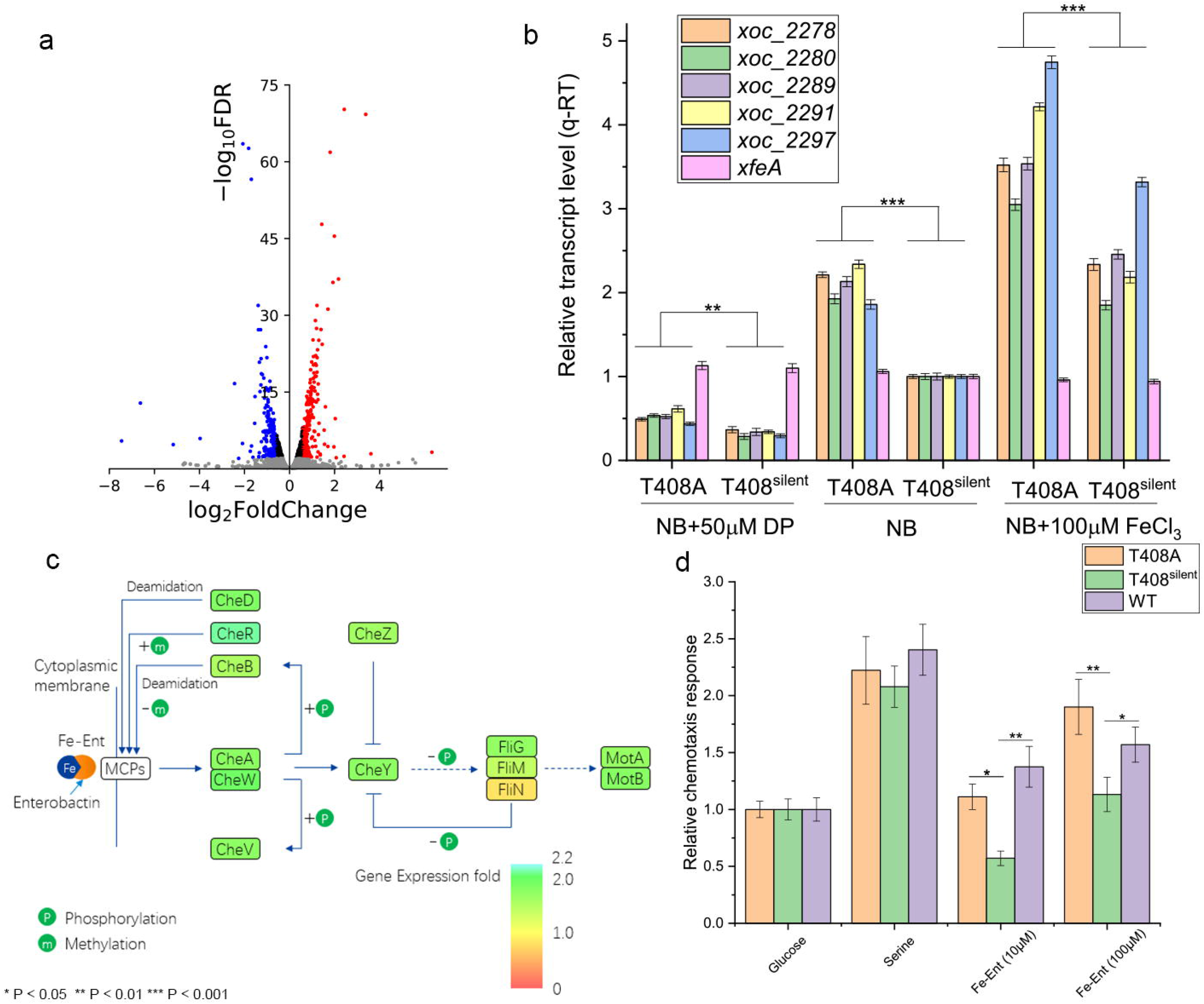
The upregulation of genes involved in chemotaxis enhances sensitivity to ferrienterobactin (Fe-Ent). (a) RNA-seq analysis of T408A and T408^silent^ strains. Volcano plot shows FDR values and fold-change of expression in T408A versus T408^silent^. Red dots indicate upregulated genes with FDR <0.01, and blue dots indicate downregulated genes with FDR <0.01. (b) Expression of selected chemotaxis pathway genes in *Xoc* strains T408A and T408^silent^. Bacterial cells were cultured in NB+50 μM DP, NB, or NB+100 μM FeCl_3_. Expression was normalized with *rpoD* and the ΔΔCT method where CT is the threshold cycle. Three independent biological replicates were carried out in this experiment. Asterisks represent significant differences between T408A and T408^silent^ at *P*<0.01 (**) or 0.001 (***) using the Student’s t-test. (c) Upregulated genes in the chemotaxis pathway based on RNA-seq. Upregulated genes are indicated in green. Model was based on the KEGG chemotaxis pathway (https://www.genome.jp/entry/map02030). (d) Chemotactic response of *Xoc* WT, T408A, and T408^silent^ strains in response to glucose (2.0 mg/mL), serine (10 mg/mL), Fe-Ent (10 μM and 100 μM) and PBS buffer (representing random diffusion). Relative chemotactic response values were calculated as a function of the number of migrated bacterial cells. Asterisks represent significant differences at *P*<0.05 (**) or 0.01 (**) using the Student’s t-test.

A capillary assay was performed to determine the chemotaxis of T408A, T408^silent^ and WT strains towards glucose (2.0 mg/mL), serine (10 mg/mL), Fe-Ent (10 μM and 100 μM) and 0.01 M pH=7.0 PBS buffer (representing random diffusion). The chemotactic response relative to glucose was calculated for each strain. The T408A strain showed a significant chemotactic response to 10 and 100 μM Fe-Ent, whereas T408^silent^ showed reduced sensitivity to Fe-Ent (Fig. 3d). There was no significant difference among strains in chemotaxis for serine.

Two MCPs, Xoc_2282 and Xoc_2291, were expressed and purified to determine whether they interact directly with FeCl_3_ or Fe-Ent using the Octet RED system (ORS) (Fig. 4a-d). The MCP/Fe-Ent association curves exceeded the 0 nm line (Fig. 4c, d), indicating that the MCPs interact directly with Fe-Ent. However, association curves of the MCP/FeCl_3_ interaction fell below the 0 nm line, which indicates lack of binding between the MCPs and Fe^3+^. The Xoc_2282/Fe-Ent and Xoc_2291/Fe-Ent interactions were also evaluated with the Biacore 8K system (Fig. 4c,d). The results showed that the MCPs directly bind Fe-Ent (Fig. 4c,d) with K_D_ values ranging from 4-6 ×10^−8^ M (Table 1). Collectively, these results indicated that T408A editing can improve the chemotactic response of bacteria for Fe-Ent, and ultimately increase intracellular iron concentrations.

**Figure 4.**
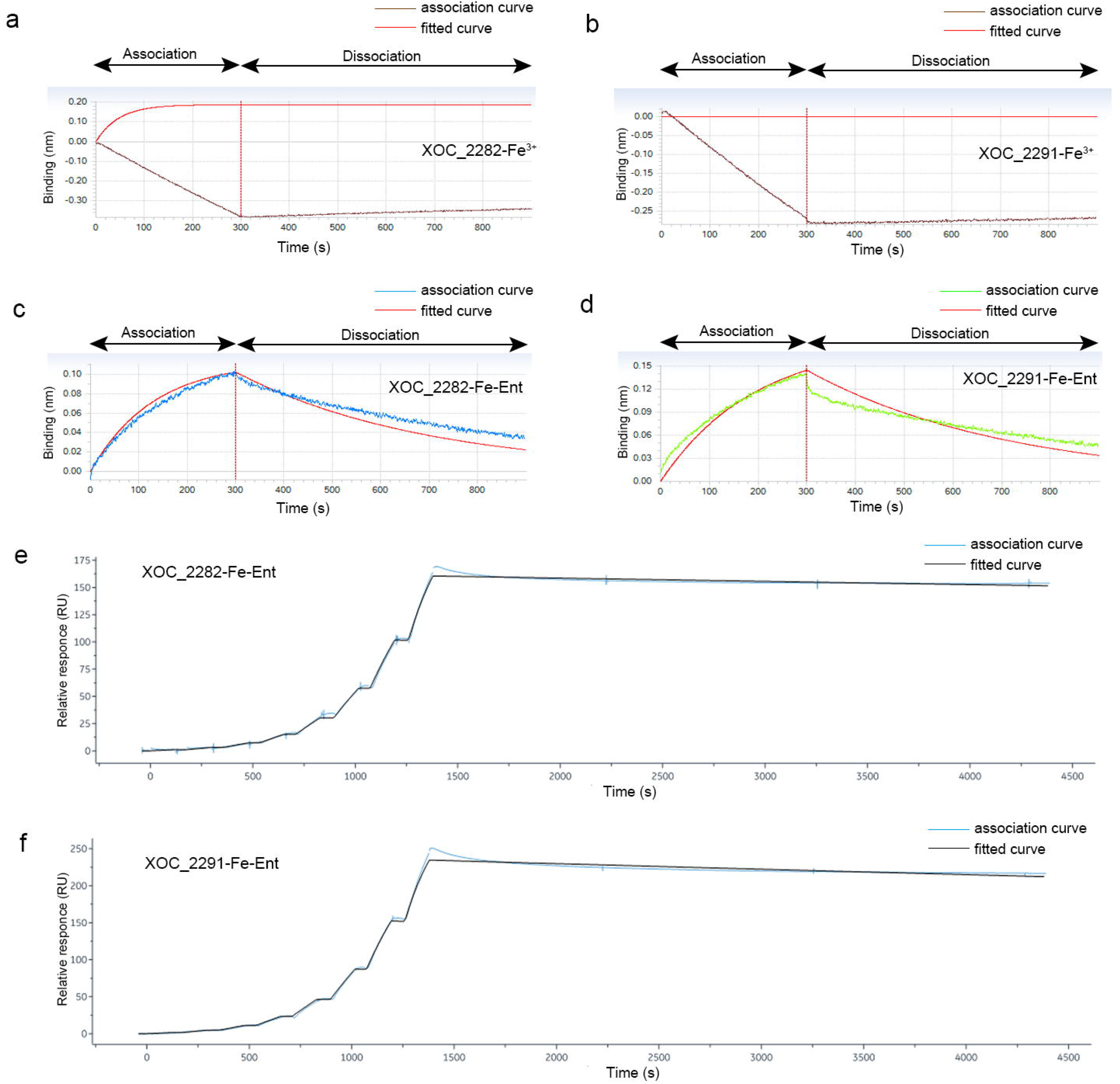
Binding analysis of MCPs Xoc_2282 and Xoc_2291 to FeCl_3_ and Fe-Ent. (a) Real-time association and dissociation analysis of (a) Xoc_2282 and (b) Xoc_2291 binding to 50 μM FeCl_3_ using the Octet Red System (ORS); brown line, association curve; red, fitted curve. Association and dissociation analysis of (c) Xoc_2282 and (d) Xoc_2291 binding to 50 μM Fe-Ent using ORS; blue and green represent association curves; red, fitted curves. The two MCPs were bound to the Ni-NTA sensor, washed and blocked. Biosensors were then incubated with FeCl_3_ or Fe-Ent in PBS buffer to facilitate association with the MCPs, and biosensors were incubated with PBS buffer to determine dissociation rates. Analysis of the (e) Xoc_2282/Fe-Ent and (f) Xoc_2291/Fe-Ent interactions with the Biacore 8K system. Eight serial two-fold concentrations of Fe-Ent were used for measuring response units (RU). PBS-T buffer was used for disassociating the Fe-Ent and MCP complex for 3000 s (blue, association curve; black, fitted curve). RUs were analyzed with Biacore Insight Evaluation software.

### T408A editing contributes to *Xoc* virulence

Leaves of six-week-old susceptible rice cv. Yuanfengzao were inoculated with *Xoc* BL256 (WT) and the T408A and T408^silent^ mutants (Fig. 5a). At 14 days post inoculation, lesions induced by the T408A mutant were 3.17□cm in length and significantly longer than lesions induced by the WT (2.33 cm) and T408^silent^ (1.88) strains (Fig.□5b). *In planta* growth assays indicated that the T408A mutant multiplied to significantly higher levels than the WT and T408^silent^ mutant (Fig. 5c). These results indicate that T408A editing enhances virulence in *Xoc*, possibly because of increased iron uptake.

**Figure 5.**
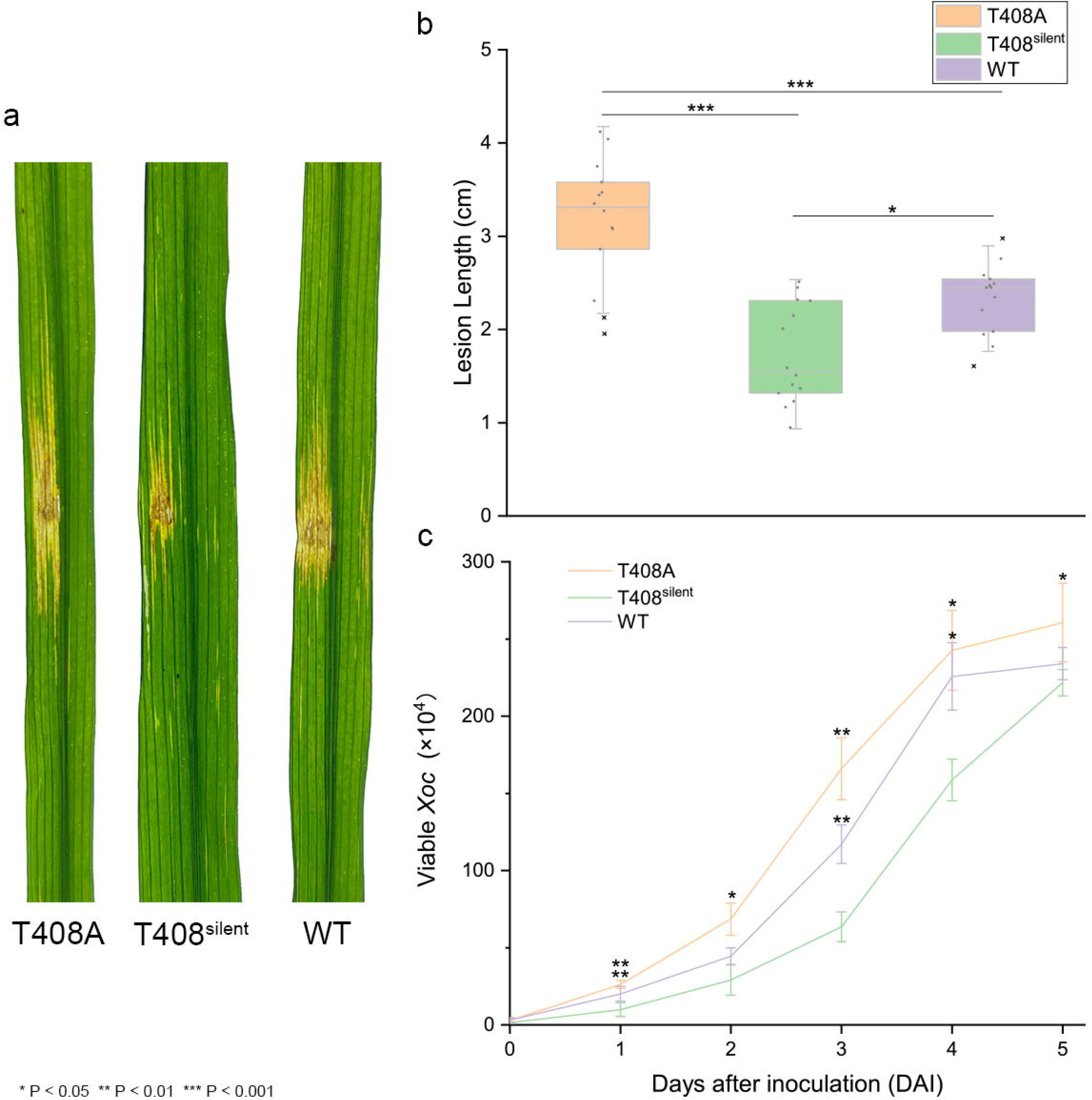
Virulence and growth of *Xoc* strains in rice cv. Yuanfengzao. Virulence was assessed by inoculating six-week-old susceptible Yuanfengzao rice plants with *Xoc* strains. (a) Symptoms on rice leaves inoculated with *Xoc* T408A, T408^silent^ and WT. (b) Lesion length of *Xoc* T408A, T408^silent^ and WT in rice cv. Yuanfengzao. Leaves (*n*=13) were inoculated with needleless syringes and evaluated for lesion length 14 days after inoculation. Values represent the mean lesion length□±□SD. The (×) indicates an abnormally high or low value that was excluded from statistical analysis. The asterisks (*) indicate significant differences between the lesion length obtained for *Xoc* T408A and WT as compared to the T408^silent^ strain (*, *P*□<□0.05; ***, *P*□<□0.001; ANOVA with Dunnett’s post-hoc correction). (c) Population dynamics of *Xoc* T408A, T408^silent^ and WT *in planta* (means ± SD). Infected leaves (*n*=3) were excised around the inoculation site, macerated, and then plated in serial dilutions to NB agar with cephalexin. *, *P* < 0.05; **, *P* < 0.01.

### Homology modeling of XfeA in *Xoc* strains

XfeA is an orthologue of FepA, the ferrienterobactin outer membrane receptor protein (Buchanan et al., 1999). FepA functions in the entry of Fe-Ent into the cell, which is an important route of iron uptake in some gram-negative bacteria (Ma et al., 2007; Newton, Igo, Scott, & Klebba, 1999). Thus, we hypothesized that T408A editing could change the efficiency of ferric siderophore entry and iron uptake. To investigate this, the secondary structure and 3D homology model of XfeA were constructed based on the crystal structure of multiple TBDR templates, including over 20 known 3D models of ferric siderophore receptors available at the Phyre2 web site (Kelley, Mezulis, Yates, Wass, & Sternberg, 2015). This approach enabled modeling of 797 residues (98%) at >90% confidence. Modeling revealed that XfeA is a 22-stranded transmembrane β-barrel protein containing an N-terminal plug domain (Fig. 6a), a configuration that is conserved in other TBDRs (Ferguson & Deisenhofer, 2002). The Thr408 residue is located on the inner side of the barrel, and replacement with the Ala residue results in a truncated β-strand (Fig. 6b, Fig. S5), thus changing the network of H bonds (Fig. 6c, 6d). Interestingly, the predicted binding region for Fe-Ent is positioned near the N-terminal plug, which is located away from the Thr408 residue (Moynié et al., 2019) (Fig. S6).

**Figure 6.**
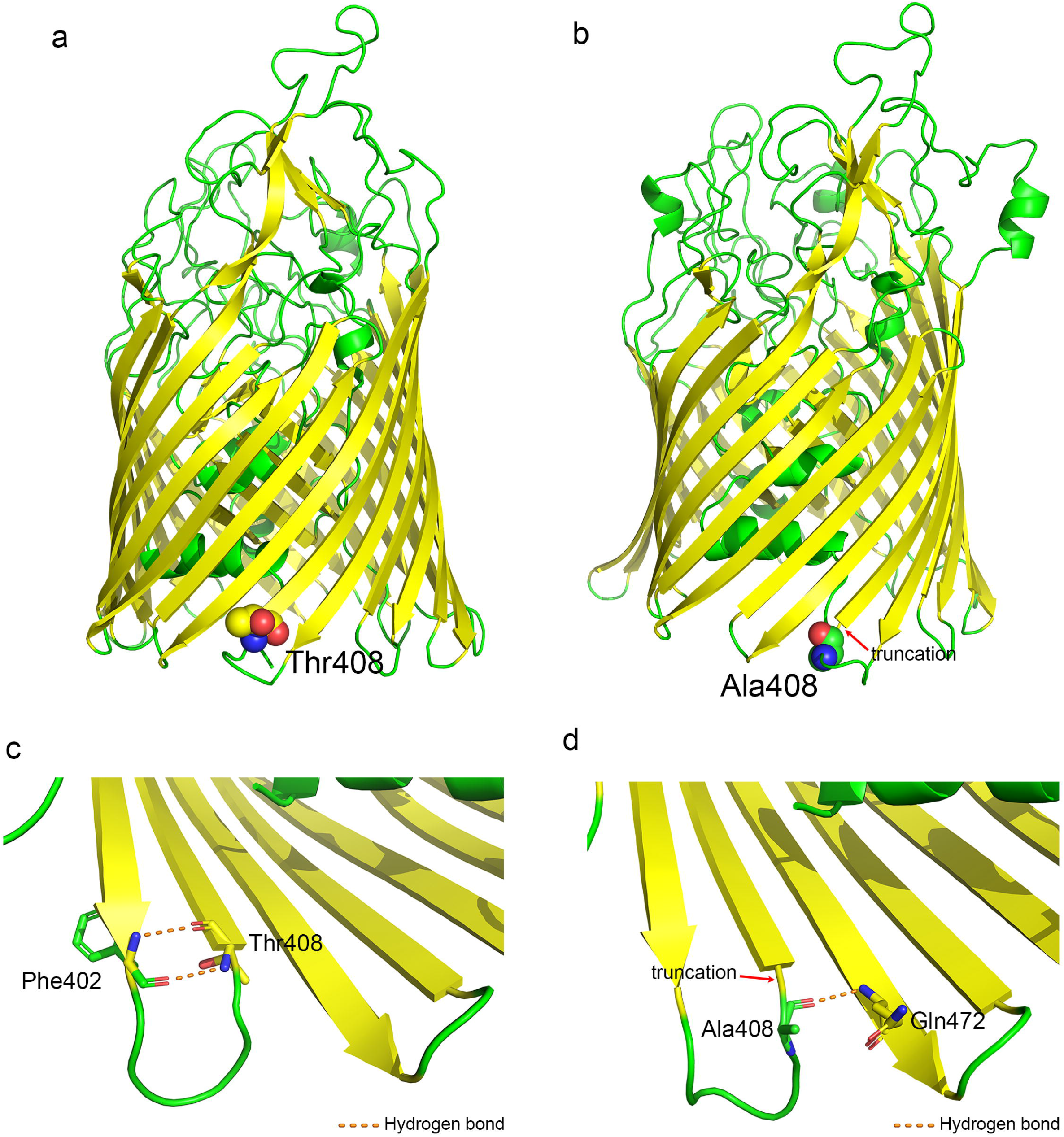
Homology modeling of XfeA in *Xoc* strains. (a) Model of XfeA in wild-type *Xoc* BL256 using multiple templates available at the Phyre2 web site. The backbone β-strands are depicted in yellow, and the 22-stranded transmembrane β-barrel and plug domains are shown in green. The threonine residue (Thr408) is depicted with spheres. (b) Model of XfeA in the T408A mutant. The red arrow indicates the truncated β-strands at the Ala408 residue, which are shown using spheres. Panels (c) and (d) show hydrogen bonding around residue 408 in XfeA. The altered H bonds (orange dotted line) surrounding Thr408 (Fig. 6c) and Ala408 (Fig. 6d) are shown. Carbon atoms at residue 408 are shown in yellow; α-helices and turns are shown in green, oxygen atoms are depicted in red, and nitrogen atoms are colored blue.

## DISCUSSION

Most bacteria deal with excessive or deficient levels of iron via the ferric uptake regulator (Fur) (Hantke, 2001), an important cytoplasmic regulator of Fe^2+^ levels in bacteria (Hassan & Troxell, 2013). Fur-mediated regulation controls the expression of many genes, including some that encode virulence factors (McHugh et al., 2003). In the current study, we show that bacteria also use post-transcriptional A-to-I editing to regulate iron uptake. In eukaryotes, A-to-I RNA editing functions in multiple regulatory processes, including splicing, microRNA targeting/processing and mRNA stability (Schaffer et al., 2020); however, A-to-I editing has only recently been described in bacteria (Bar-Yaacov et al., 2017; Nie et al., 2020; Safra et al., 2017).

In this study, *Xoc* responded to iron-limiting conditions by A-to-I RNA editing in *xfeA*. A mutant strain was generated with fixed A-to-I editing in *xfeA* and designated T408A. Strain T408A showed improved growth in iron-limiting conditions and a significant chemotactic response to ferrienterobactin. A-to-I RNA editing in *xfeA* also increased expression of multiple genes in the chemotaxis pathway, including two methyl-accepting chemotaxis proteins, Xoc_2282 and Xoc_2291. Interestingly, both Xoc_2282 and Xoc_2291 interacted with Fe-Ent but not FeCl_3_. It is tempting to speculate that XfeA functions analogous to FepA, the ferrienterobactin outer membrane receptor protein that facilitates the transport of Fe-Ent into the cell (Ma et al., 2007; Newton et al., 1999).

MCPs are the predominant chemoreceptors in bacteria and regulate diverse cellular activities (Ud-Din & Roujeinikova, 2017). A typical MCP contains a domain that interacts directly with the ligand, which then transduces the signal to downstream genes (Milburn et al., 1991; Muok et al., 2019). In this study, MCPs Xoc_2282 and Xoc_2291 interacted with Fe-Ent, and binding kinetic constants were calculated. The interaction of MCPs with the ligand, Fe-Ent, was associated with increased activity of CheA and other chemotaxis-related genes (Fig. 3c, Table S3); this is consistent with activation of the chemotaxis signal transduction pathway. In *E. coli*, a MCP-CheW-CheA complex transduces signals to the response regulator CheY by phosphorylation (Wuichet & Zhulin, 2010). A similar phenomenon might occur in *Xoc* and cause upregulation of additional chemotaxis genes in response to low iron availability and A-to-I editing in XfeA.

Although the MCPs Xoc_2282 and Xoc_2291 interacted with Fe-Ent, *Xoc* is not known to synthesize enterobactin as a siderophore. The predominant siderophore produced by *Xanthomonas* spp. is xanthoferrin, an α-hydroxycarboxylate molecule that is known to be synthesized by *X. oryzae* (Pandey & Sonti, 2010). Interestingly, *Xoc* does contain several genes associated with enterobactin synthesis; e.g. *xoc_2573* (phosphopantetheinyl transferase, EntD), *xoc_2574* (Ent synthase subunit F) and *xoc_2575* (ATP-dependent serine activating enzyme, EntF). Although these three genes were differentially expressed in the RNA-seq analysis of *Xoc* strains T408A and T408^silent^ (differential expression data, https://drive.google.com/file/d/1FTiS4tQpVsyHcSvJKk4NtZZludVntRxE/view?usp=sharing), we were unable to conclusively demonstrate enterobactin synthesis in *Xoc*. It remains possible that *Xoc* synthesizes an enterobactin-like analogue; however, another possibility is that *Xoc* does not synthesize an Ent-like analogue, but instead retains the FepA-D proteins for exogenous ferrienterobactin uptake. For example, a recent study with *X. oryzae* pv. *oryzae* indicated that the pathogen produces several FecA and Ent-like receptors *in planta*, thus allowing the pathogen to acquire iron from heterologous ferric siderophores (González et al., 2012). In this respect, *X. oryzae* might resemble *Vibrio cholera* where siderophore piracy has been established (Byun, Jung, Chen, Valencia, & Zhu, 2020; Wyckoff, Allred, Raymond, & Payne, 2015).

The detection of iron concentrations by pathogenic bacteria is a critical factor in survival and establishment of a successful infection (L. Wang et al., 2016). The TonB complex provides the energy required for active transport of ferric siderophores through TonB-dependent transporters (TBDTs) that are located in the outer membrane of gram-negative bacteria (Noinaj, Guillier, Barnard, & Buchanan, 2010). In *X. oryzae*, ferric enterobactin outer membrane receptors have been reported as potential virulence factors, partly due to their upregulation in rice (Carnielli, Artier, de Oliveira, & Novo-Mansur, 2017; González et al., 2012; Xu et al., 2015). In this study, we show that the *Xoc* T408A mutant is more virulent than the WT, and this is attributed to the fixed A-to-I editing in the TonB-dependent receptor XfeA and increased iron uptake. Our findings further support the importance of iron, siderophore uptake, and TonB-dependent receptors in the virulence of *Xanthomonas* spp. (Timilsina et al., 2020).

A curious finding in the present study was the TadA-independence of A-to-I editing in *xfeA*; this was unexpected due to the presence of motifs and structures in *xfeA* that would be recognized by TadA. There are reports indicating duplication of ancestral *tadA* in various bacteria (Torres et al., 2014); however, database searches for other *tadA* homologues in the BLS256 genome were unsuccessful. Since the A-to-I editing site in *xfeA* is largely unaffected in the *tadA* deletion mutant, we suggested that the unknown enzyme might be a specialized mRNA deaminase unlike TadA. The editing of mRNAs by adenosine deaminases acting on RNA (ADARs) is conserved in metazoans; however, recent work with filamentous fungi lacking ADAR orthologues has demonstrated that other mechanisms for A-to-I RNA editing exist (Bian, Ni, Xu, & Liu, 2019). It is also important to note that a new group of A-to-I RNA editing enzymes was recently described in *E. coli* and named *r*estriction by an *a*denosine *d*eaminase *a*cting on *R*NA (RADAR) (Gao et al., 2020). Although we were unable to identify obvious homologues for RADAR genes in *Xoc*, the discovery of this group of proteins further illustrates the capacity of bacteria to develop TadA-independent mechanisms for RNA editing.

Since both excessive and deficient levels of iron can be harmful, bacteria use global regulators such as Fur to detect iron and regulate the expression of genes involved in iron storage, uptake, and efflux (Bradley et al., 2020). Our study demonstrates that *Xoc* also uses A-to-I editing in *xfeA* to modulate iron acquisition via ferric siderophore transport across the outer membrane (Fig. 7). A-to-I editing in *xfeA* increases when iron is limiting and causes changes in the hydrogen-bonding network of XfeA; this facilitates transport of the ferric siderophore through the XfeA channel. A-to-I RNA editing in *xfeA* leads to increased expression of genes encoding MCPs, which interact with the ferric siderophore in the cytoplasmic membrane. Genes in the chemotaxis pathway are also induced as a result of A-to-I editing, and *Xoc* is chemotactically attracted to the ferric siderophore in the external milieu. When iron is plentiful, A-to-I editing in *xfeA* decreases, and less ferric siderophore traverses the XfeA channel of (Fig. 7).

**Figure 7.**
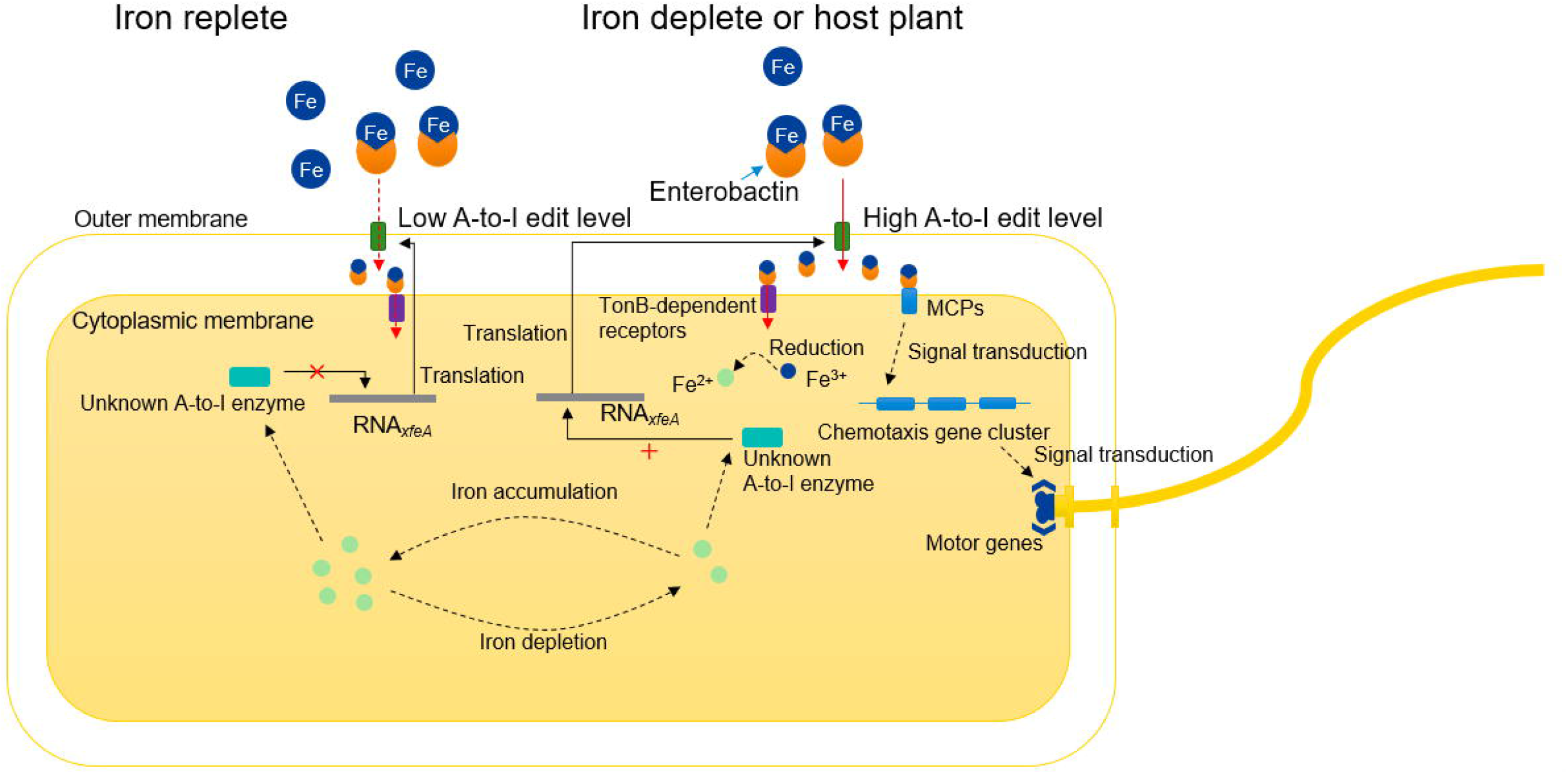
Proposed regulation of iron homeostasis by A-to-I editing in *xfeA*. Under iron-depleted conditions or *in planta*, T408A editing levels within *xfeA* are enhanced and genes encoding MCPs are upregulated. *X. oryzae* is chemotactically attracted to Fe-Ent, which is transported across the bacterial membranes. Fe^3+^ is converted to Fe^2+^ in the cytoplasm and used for various reactions, including responses and pathways that contribute to pathogen virulence. In iron replete conditions, A-to-I editing in *xfeA* is reduced.

In a recent study, we demonstrated that an A-to-I editing event in *fliC*, which encodes a flagella filament protein in *Xoc*, occurred in response to oxidative stress (Nie et al., 2020). It is important to note that *Xoc* is a seed-borne pathogen that is exposed to both oxidative stress and changing iron conditions in the host plant. The current study expands our knowledge of RNA editing in pathogenic bacteria and provides a mechanism for adapting to iron-deficient conditions. A-to-I RNA editing provides the bacterial cell a quick and rapid way to recode protein products that are appropriate for changes that occur during pathogenesis. It will be interesting to see if other pathogens use a similar mechanism to adapt to fluctuations in iron levels.

## MATERIALS AND METHODS

### Bacterial strains, plasmids, plant materials, and reagents

The bacterial strains and plasmids used in this study are described in Table S1. *Xoc* strain T408A is a derivative of BLS256 containing a mutation (T<u>A</u>CG to T<u>G</u>CG) that changes the threonine at amino acid residue 408 in XfeA to alanine. T408^silent^ contains a synonymous mutation (TAC<u>G</u> to TAC<u>A</u>) in *xfeA* that blocks post-transcriptional A-to-I editing at amino acid 408. *E. coli* strains BL21-2282 and BL21-2291 harbor pET-30a::*xoc_2282* and pET-30a::*xoc_2291*, respectively, and were used to produce the MCPs Xoc_2282 and Xoc_2291.

*E. coli* strains were cultured in Luria-Bertani (LB) medium at 37°C (Maniatis, Fritsch, & Sambrook, 1982). *Xoc* BLS256 strains were cultured at 28°C in nutrient broth (NB) or nutrient agar (NA) as described previously (Nie et al., 2020). The final concentrations of antibiotics in μg/mL were as follows: kanamycin, 25; streptonigrin, 1; and cephalexin, 40. Filter-sterilized 2,2’-dipyridyl (AR, Sinopharm Chemical Reagent Co., Ltd, China) and FeCl_3_ (Sinopharm Chemical Reagent Co.) were prepared as 10 mM stocks and diluted to 50-150 μM with NB when used. A crude extract containing enterobactin was kindly provided by Prof. Fu-Zhou Xu of Beijing Academy of Agriculture and Forestry Sciences. Enterobactin was purified as described previously (Zeng, Xu, & Lin, 2009) and diluted in H_2_O to prepare a 2 mM solution; FeCl_3_ (2 mM) was then added to prepare the Fe-Ent stock solution (1 mM).

Rice seeds were obtained from the International Rice Research Institute and grown in a greenhouse as described previously (Nie et al., 2020)

### Bacterial mutant and strain construction

Mutant strains were constructed as described previously (S. Wang et al., 2020). The coding region of *xfeA* was first amplified from *Xoc* BLS256 with primers *xfeA* F/R (Table S2), digested with *Sal*I/*Xho*I, and subcloned in pKMS1 (Xie et al., 2011). The Fast Mutagenesis System (Transgen Biotech, Beijing, China) was used to obtain clones with the T408A and T408^silent^ point mutations. Primers T408A F/R and T408^silent^ F/R were used to introduce point mutations into *xfeA* by PCR (Table S2), and mutated *xfeA* alleles were introduced into BL256 by double homologous recombination as described previously (S. Wang et al., 2020).

The methyl-accepting chemotaxis proteins, Xoc_2282 and Xoc_2291, were overexpressed in *E. coli* BL21(DE3). Full-length fragments of *xoc_2282* and *xoc_2291* were amplified with *Pfu* polymerase (TransGen Biotech, Beijing, China) using the xoc2282 and xoc2291 F/R primers (Table S2). These fragments were cloned, digested with *Bam*HI*/Hin*dIII, ligated into pET-30a (+), and then transformed into *E. coli* BL21 (DE3) by heat shock at 42□ for 45 s. Transformants were selected on LB agar with kanamycin.

### RNA secondary structure prediction

A sequence extending −25 to +25 bp from the edited site was used to model RNA secondary structure using the RNAfold web server (http://rna.tbi.univie.ac.at/cgi-bin/RNAWebSuite/RNAfold.cgi) as described (S. Wang et al., 2020).

### Protein structure prediction

For secondary structure and 3D modeling, the XfeA sequence (Thr408 and Ala408) was submitted to the Phyre2 web site (http://www.sbg.bio.ic.ac.uk/phyre2) for modeling with multiple templates (Kelley et al., 2015); exactly 797 residues (98%) were modeled at >90% confidence. Based on the XfeA models, the Thr408 and Ala408 residues were modeled using spheres where carbons in β-strands were colored yellow and other carbons were shown in green. The hydrogen bonds located near Thr408 and Ala408 were displayed with Pymol (Schrödinger, 2015). The model of ferric enterobactin was retrieved from the PubChem website (CID: 34231), and the interaction with XfeA was predicted by Autodock Vina (Trott & Olson, 2010). Default values were used for the iteration limit and RMS gradient test.

### Bacterial growth assays in response to iron depletion

Optical density (OD) was measured with a Bioscreen C instrument (Labsystem, Helsinki, Finland) as described previously (Nie et al., 2020). OD values at 420-580 nm were measured at 15 min intervals for 24 h with continuous shaking at 28°C, and experiments included four independent replicates. Pairwise comparisons of growth curves for strains and growth conditions were analyzed with the *F*-test and compared with the curve obtained for T408^silent^ mutant. OriginPro v. 9.5.1.195 was used to graph, display and analyze the data.

### Analysis of intracellular iron

*Xoc* cells were analyzed for iron content by ICP-OES (Optima 8000, PerkinElmer, MA USA) as reported previously (L. Wang et al., 2016). *Xoc* cells were grown to an OD_600_=1.0 in NB, collected by centrifugation, washed three times in sterile PBS (NaCl 8.5g/L, Na_2_HPO_4_ 2.2g/L, NaH_2_PO_4_ 0.4g/L, pH=7.0), and then inoculated into two-fold volumes of fresh NB supplemented with 50 μM DP or 100 μM FeCl_3_. After 3-5 h (OD_600_~0.6), cells were harvested by centrifugation and washed in PBS; pelleted cells were then dried at 65□ for 48 h, and digested with acid (HNO_3_–HClO_4_, 4:1, v/v). The digested cellular material was transferred to a 25 mL volumetric flask, diluted to 25 mL in deionized water, and iron was measured by ICP-OES. Samples that were not digested in acid were analyzed in parallel as controls. Iron concentrations were calculated by dividing the iron atomic value for 10^9^ cells.

### Streptonigrin survival assays

Survival in response to streptonigrin exposure was evaluated as described previously with slight modifications (Si et al., 2017). Log-phase (OD_600_=0.6) *Xoc* cells were harvested by centrifugation, washed three times in 0.01 M PBS buffer, and exposed to 1 μg/mL streptonigrin for 16 h at 28°C. Cultures were then diluted, inoculated to NA, and colonies were counted as described (Nie et al., 2020). Survival was calculated by comparing viable cells counts with and without streptonigrin. This assay was performed in triplicate.

### Exposure of *Xoc* strains to DP or FeCl_3_

*Xoc* BLS256 was incubated at 28□ to OD_600_=1.0 in NB with DP (50, 100 or 150 μM) or 100 μM FeCl_3_. Aliquots were removed and cells were harvested by centrifugation at 4°C. Pellets were washed twice in cold PBS, and total RNA was extracted with the RNeasy Protect Bacteria Mini Kit (Qiagen). This experiment contained two biological replicates.

### PCR and qRT-PCR

Total RNA (10 μg/sample) was isolated and used to synthesize cDNA as described previously (Nie et al., 2020). The *xfeA* transcript in cDNA samples was sequenced using the c T408 F/R primers (Table S2). The sequencing chromatograms were analyzed with Chromas Lite (Technelysium, Brisbane, Australia), and the frequency of editing was estimated by ratiometric A/G measurement as described (Nie et al., 2020).

The EasyPure RNA Kit was used to purify RNA as recommended (Transgen Biotech), and 1 μg of RNA was used to synthesize cDNA with the Magic 1st cDNA Synthesis Kit (Magic Biotech, Hangzhou, China). The cDNA product (20 μl) was diluted to 100 μl and used for qRT-PCR with Magic SYBR Green qPCR Mix (Magic Biotech) and the ABI 7500 quantitative PCR system (Applied Biosystems, Foster City, CA). Expression was normalized with *rpoD* using the ΔΔCT method as described (Nie et al., 2020). Experiments included three independent biological replicates.

### RNA sequencing and RNA-seq data analysis

RNA-seq libraries were prepared using the Illumina Paired End Sample Prep kit as described previously (Fang et al., 2019). After removal of adaptors and low quality reads, RNA-seq reads were aligned to the *Xoc* BLS256 genome using Tophat 2.0.7 (Trapnell, Pachter, & Salzberg, 2009), allowing for a maximum of two mismatched nucleotides. If reads mapped to more than one location in the genome, only the site showing the highest score was retained. Reads that mapped to tRNA or rRNA regions were removed; the remaining reads were mapped to the genome with HISAT2 (Kim, Paggi, Park, Bennett, & Salzberg, 2019) and used to generate a volcano plot. Bioconductor package edgeR (Robinson, McCarthy, & Smyth, 2010) with TMM normalization was used to determine DEGs as described (Nie et al., 2020). Reproducibility was evaluated for two replicate experiments using pairwise linear correlation analysis prior to comparing RNA-seq profiles.

DEGs with significance (FDR < 0.01; fold change > 2) were selected for further analysis (differential expression data, https://drive.google.com/file/d/1FTiS4tQpVsyHcSvJKk4NtZZludVntRxE/view?usp=sharing). Treeview 1.1.6 and Cluster 3.0 (de Hoon, Imoto, Nolan, & Miyano, 2004; Saldanha, 2004) were utilized to produce heatmaps based on reads/kb of transcript per million mapped reads (RPKM) (de Hoon et al., 2004; Saldanha, 2004).

### Capillary chemotaxis assay

Chemotaxis was evaluated with the capillary method as described previously (Kumar Verma, Samal, & Chatterjee, 2018) with slight modifications. *Xoc* strains were grown in NB to OD_600_=1.0, centrifuged at 800 ×g for 6 min, washed with PBS three times and resuspended in 1 ml of PBS. Sterilized capillary tubes containing filter-sterilized glucose (attractant control, 2.0 mg/mL), serine (10 mg/mL), Fe-Ent (10 and 100 μM) or 0.01 M PBS buffer (negative control) were incubated with *Xoc* strains at 28 °C for 4 h. To determine bacterial cell counts, the contents of capillaries were serially diluted (10-fold) in PBS and plated to NA (Fig. S3). The chemotaxis response relative to glucose was calculated as the number of migrated bacterial cells in the capillary minus the cells counted in PBS buffer, which would be attributed to random motility or diffusion.

### Kinetic analysis of MCP binding to FeCl_3_ and Fe-Ent

The MCPs Xoc_2282 and Xoc_2291 were purified from *E. coli* BL21-xoc2282 and BL21-xoc2291 (Table S1) using the BeyoGold His-tag purification resin (Beyotime, Shanghai, China); proteins were then concentrated with a molecular filter (30 kDa cut-off, (Millipore China, Shanghai, China).

Kinetic analysis of binding was performed with the Octet Red 96 system (ForteBio, Fremont, CA, USA) as described previously (Li et al., 2015) with several modifications. FeCl_3_ or Fe-Ent was dissolved in PBS at a concentration of 50 μM, and the MCPs Xoc_2282 and Xoc_2291 were diluted in PBS at 10 μM. After a baseline step, His-tag labeled Xoc_2282 and Xoc_2291 were bound to the Octet Red Ni-NTA biosensors, washed and blocked with BSA buffer (0.2% BSA and 0.02% Tween-20 in PBS buffer). The biosensors were incubated for 300 s with 50 μM FeCl_3_ or Fe-Ent to facilitate association and incubated with PBS for 600 s to determine the dissociation rate. Octet Data Analysis HT v. 12 software was used to fit the curve.

For analysis with the Biacore 8K surface plasmon resonance system (Jason-Moller, Murphy, & Bruno, 2006), Xoc_2282 or Xoc_2291 (10 μg/mL) were dissolved in sodium acetate buffer (pH=4.0) and bound to the CM5 chip for 1200 s in PBS buffer (pH=7.4) to ensure that >10,000 units of protein were loaded. Eight, serial two-fold concentrations of Fe-Ent were used for measuring and recording the response units (RU), and the final concentration of Fe-Ent was 0.078, 0.156, 0.3125, 0.625, 1.25, 2.5, 5 and 10 μM. PBS-T buffer (0.01 M PBS, 0.05% Tween 20, pH=7.4) was used to disassociate Fe-Ent and MCP proteins for 3000 s. Response units were analyzed with Biacore Insight Evaluation software and kinetic rate constants were calculated (K_D_=K_d_/K_a_; where K_D_ is the dissociation equilibrium constant, K_d_ is the dissociation constant, and K_a_ is association constant).

### Plant inoculation assays

Suspensions of *Xoc* (OD_600_=0.6) were used to inoculate six-week-old rice. Symptoms and bacterial populations were described and enumerated using established methods (Nie et al., 2020).

### Data availability

All sequence data generated in this study were deposited in NCBI as BioProject no. PRJNA673071.

## ACKNOWLEDGEMENTS

We thank Prof. Fu-Zhou Xu for providing Fe-Ent and Prof. Wei Qian for valuable suggestions. This work was supported by National Key R&D Program of China (2018YFD0201202, 2017YFD0201108), the Agri-X Interdisciplinary Fund of Shanghai Jiao Tong University (Agri-X2017010), the State Key Laboratory for Biology of Plant Diseases and Insect Pests (SKLOF201802), Shanghai Committee of Science and Technology (19390743300 and 21ZR1435500), National Natural Science Foundation of China (31200003, 31770772) and Joint Research Funds for Translational Medicine at Shanghai Jiao Tong University (ZH2018ZDA06).

## Appendixes

**Figure S1.** Normalized expression levels of selected chemotaxis pathway genes (*xoc_2278*, *xoc_2280*, xoc_2289, xoc_2291, and xoc_2297) and *xfeA* in *Xoc* BLS256. Wild-type BL256 was cultivated in NB, NB+50 μM 2,2’-dipyridyl (DP, iron chelator), and NB+100 μM FeCl_3_. Gene expression levels were calculated relative to *rpoD* using the ΔΔCT method, where CT is the threshold cycle. Three independent biological replicates were carried out in this study.

**Figure S2.** Prediction of the mutation site in *Xoc* T408A. RNA secondary structure analysis (http://rna.tbi.univie.ac.at/) showed that the edited site was embedded within a loop (see arrow). Color is used to show base-pair probabilities.

**Figure S3.** Diagram of capillary chemotaxis assay.

**Figure S4.** Verification of *xfeA* T408A A-to-I editing site by Sanger sequencing using gDNA as the template. Chromatograms show editing in *Xoc* BL256 (wild-type, WT) grown in NB, NB+50 μM DP, NB+100 μM DP, NB+150 μM DP, and NB+100 μM FeCl_3_. The *Xoc ΔtadA* and T408^silent^ mutants were grown in NB and included for comparison.

**Figure S5.** Predicted secondary structure of XfeA based on analysis with the Phyre2 web site. Structure of XfeA in *Xoc* (a) T408^silent^ (no editing) and (b) the T408A mutant (with editing). The latter mutant contains a truncation in the β-strand (red arrow and rectangle).

**Figure S6.** Homology modeling of XfeA showing the predicted site of ferrienterobactin binding. The model was constructed with AutoDock Vina.

Table S1. Strains and plasmids used in this study.

Table S2. Primers used in this study.

Table S3. Differentially regulated chemotaxis genes in *Xoc* T408A and T408^silent^ identified by RNA-seq.

Table S4. A-to-I RNA editing in *Xoc* T408A strain and T408^silent^ strain.

## Notes

### Competing Interest Statement

The authors have declared no competing interest.

