## Supplementary material for "A-to-I mRNA editing in a ferric siderophore receptor improves competition for iron in *Xanthomonas oryzae*": Table S1 Strains and plasmids.docx

Table S1 Strains and plasmids used in this study.

| Strain or plasmid | Relevant characteristics | Source or reference |
| --- | --- | --- |
| Strains | | |
| *Escherichia coli* | | |
| DH5α | F^-^ Φ80d *lacZ*ΔM15Δ(lacZYA-argF) U169 *recA1 endA1, hsdR17*(r_k_^-^,m_k_^+^) *phoA supE44*λ^-^ *thi-1 gyrA96 relA1* |  |
| BL21(DE3) | F^-^ *ompT* *hsdS*(r_B_^-^ m_B_^-^) *gal* *dcm* (DE3) | Transgen Biotech, Beijing, China |
| BL21-xoc2282 | BL21(DE3) harboring pET-30a::*xoc_2282*; Km^r^ | This study |
| BL21-xoc2291 | BL21(DE3) harboring pET-30a::*xoc_2291*; Km^r^ | This study |
| *Xanthomonas oryzae* pv. *oryzicola* | | |
| BLS256 | Wild-type | (Bogdanove et al., 2011) |
| T408A | BLS256 containing a mutation (ACG to GCG) that changes Thr to Ala at residue 408 in XfeA | This study |
| T408^silent^ | BLS256 containing a synonymous mutation (ACG to ACA) in *xfeA* that blocks A-to-I RNA editing at amino acid 408 | This study |
| Plasmids | | |
| pKMS1 | Km^R^; R6K-based suicide vector; requires the *pir*-encoded π protein for replication | (Li et al., 2011) |
| pET-30a(+) | Km^R^; contains N-terminal His-/thrombin/S-tag/enterokinase and C-terminal His-tag sequence; expression controlled by T7 RNA polymerase | Novagen, Madison, WI, USA |
| pKMS1::xfeA | Fragments encoding *xfeA* were amplified from *Xoc* BLS256 with primers *xfeA* F/R (Table S2), digested with *Sal*I/*Xho*I, and subcloned into the *Sal*I/*Xho*I site of pKMS1 | This study |
| pKMS1::T408A | Contains A to G point mutation in *xfeA* that changes amino acid residue 408 from Thr to Ala; cloned in pKMS1 for homologous recombination, Km^R^ | This study |
| pKMS1::T408^silent^ | Contains G to A point mutation in *xfeA* that blocks A-to-I RNA editing in residue 408; cloned in pKMS1 for homologous recombination, Km^R^ | This study |
| pET-30a::*xoc_2282* | Contains a 2382 bp *Bam*HI/*Hin*dIII fragment encoding *xoc_2282* in pET-30a(+), Km^R^ | This study |
| pET-30a::*xoc_2291* | Contains a 2109 bp *Bam*HI/*Hin*dIII fragment encoding *xoc_2291* in pET-30a(+), Km^R^ | This study |

^a^ Km^R^, kanamycin resistance.
