## Supplementary material for "A-to-I mRNA editing in a ferric siderophore receptor improves competition for iron in *Xanthomonas oryzae*": Table S2 Primers.docx

Table S2. Primers used in this study

| Application | | Primers^a^ | Sequences (5’ to 3’)^a,b^ |
| --- | --- | --- | --- |
| T408A point mutation | Amplification of *xfeA* | *xfeA* F (Sa) | GCGTCGACACAGCAAGATCGGCAAGG |
|  |  | *xfeA* R (Xh) | CCCTCGAGGGTAGTTGTCCAGGATGTTCT |
|  | Generation of T408A point mutation in *xfeA* | T408A F | TCGACGCCGTTGGTGCTGCGCACACCTTCG |
|  |  | T408A R | CAGCACCAACGGCGTCGAACACTCGGGTCA |
| T408^silent^ | Generation of T408 point synonymous mutation in *xfeA* | T408Silent F | GACGCCGTTGGTGCTACACACACCTTCGGCA |
|  |  | T408Silent R | TGTAGCACCAACGGCGTCGAACACTCGGGT |
| Amplification of *xfeA* cDNA including T408 point mutation | | c T408 F | CGCAAACCGTGGGCAATT |
|  |  | c T408 R | CCAGATCAGTGGAGAACTTGTC |
| Amplification of *xoc_2282* | | *xoc_2282* F (H) | CCAAGCTTGAACTCTTGCCAGTGACCGTC |
|  |  | *xoc_2282* R (B) | CGGGATCCATGCAATGGATCAACAATCTG |
| Amplification of *xoc_2291* | | *xoc_2291* F (H) | CCAAGCTTGAACTCCTGCCAGCTGGTC |
|  |  | *xoc_2291* R (B) | CGGGATCCATGAACGACCATACCTATCAG |
| qPCR analysis of *xfeA* | | q *xfeA* F | CCTGGCTCAACACCAAGATT |
|  |  | q *xfeA* R | GCATCGTAATCCCAGTTGCC |
| qPCR analysis of *xoc_2278* | | q *xoc_2278* F | CGATGGCGATCAGACGAT |
|  |  | q *xoc_2278* R | GTCAGCAAGGTCAAGATGG |
| qPCR analysis of *xoc_2280* | | q *xoc_2280* F | ATCACATTGCGACAGAACA |
|  |  | q *xoc_2280* R | CAAGTGCCGAGTCATTCC |
| qPCR analysis of *xoc_2289* | | q *xoc_2289* F | GTAGATCAGCGTGGACAC |
|  |  | q *xoc_2289* R | CTTGAGCGAACAGACCTT |
| qPCR analysis of *xoc_2291* | | q *xoc_2291* F | GAGGCTTTCCACCATCAC |
|  |  | q *xoc_2291* R | CCTGTCTTCGGTCAATCG |
| qPCR analysis of *xoc_2297* | | q *xoc_2297* F | ATCTGCTCTGCTTGGTAG |
|  |  | q *xoc_2297* R | GCGTATCGTCTGATATTCG |
| qPCR analysis of *rpoD* (internal control) | | *rpoD* F | CGACAACACCACCAACATCAATC |
|  |  | *rpoD* R | GCTTACCGACCTCTTCCAACG |

^a^ The following restriction sites were introduced into primer sequences: B, *Bam*HI; H, *Hin*dIII; Sa, *Sal*I; and Xh, *Xho*I. Restriction sites are underscored in the primer sequences.

^b^ Nucleotides in red font indicate introduction of a point mutation.
