## Supplementary figures and images for "A-to-I mRNA editing in a ferric siderophore receptor improves competition for iron in *Xanthomonas oryzae*"

### fig s1.tif

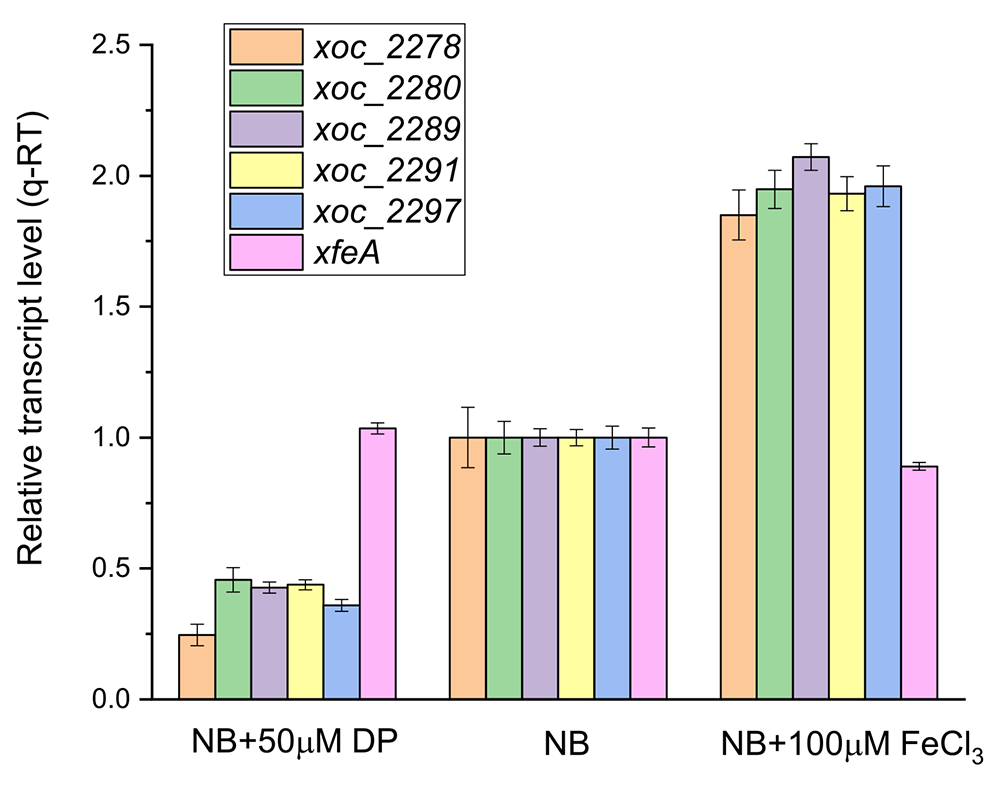

### fig s2.tif

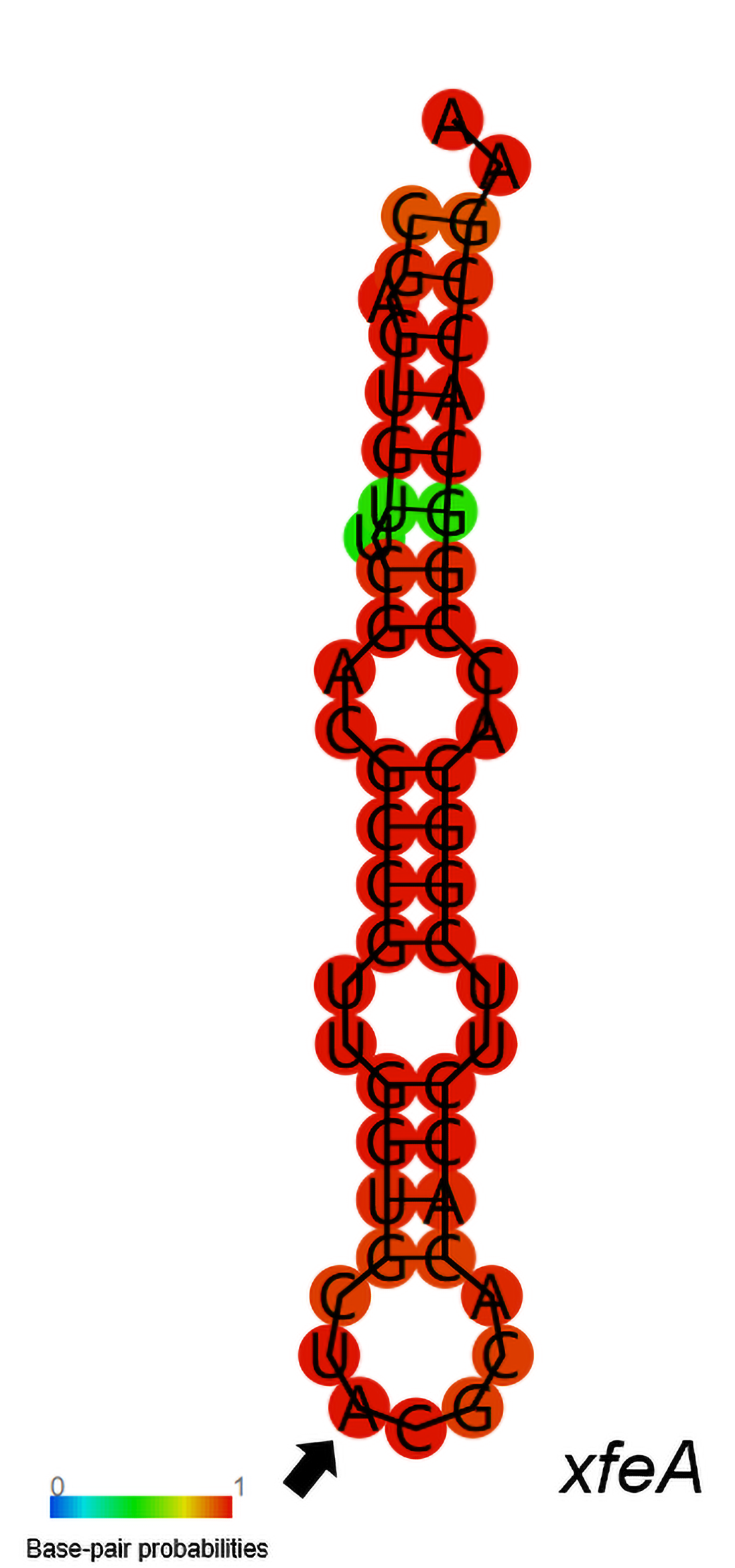

### fig s3.tif

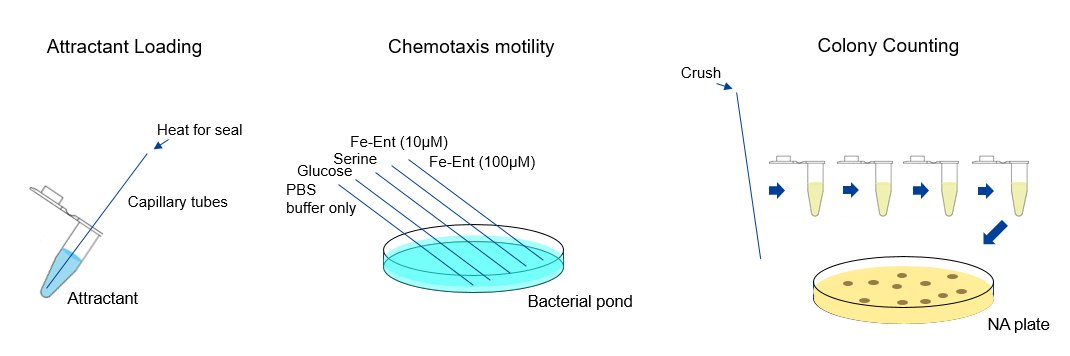

### fig s4.tif

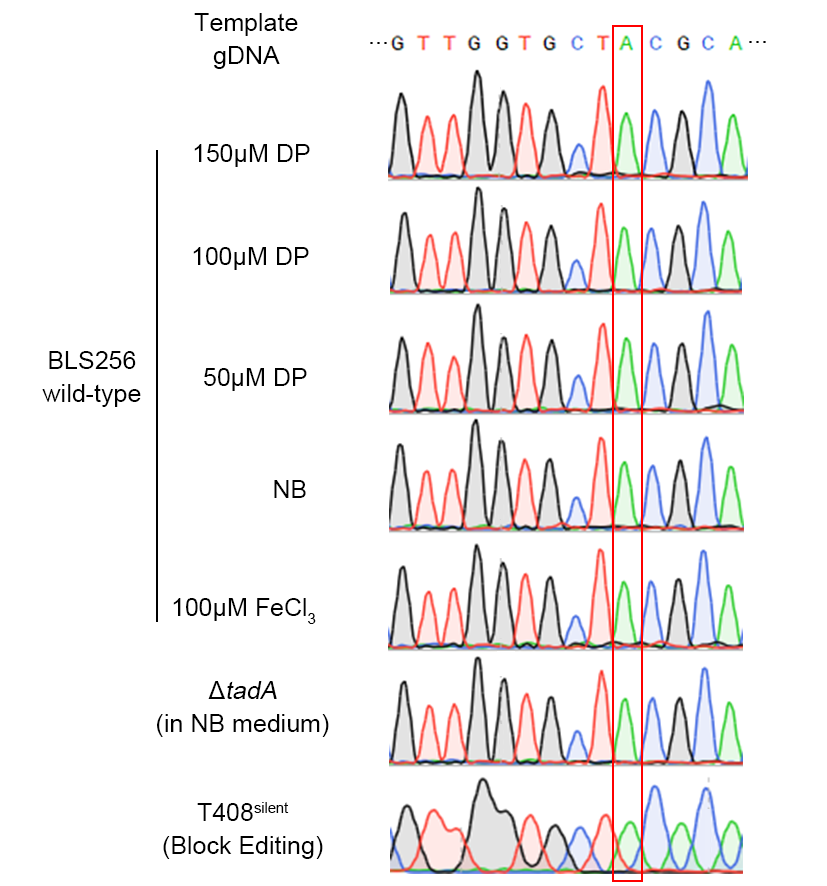

### Fig S5.tif

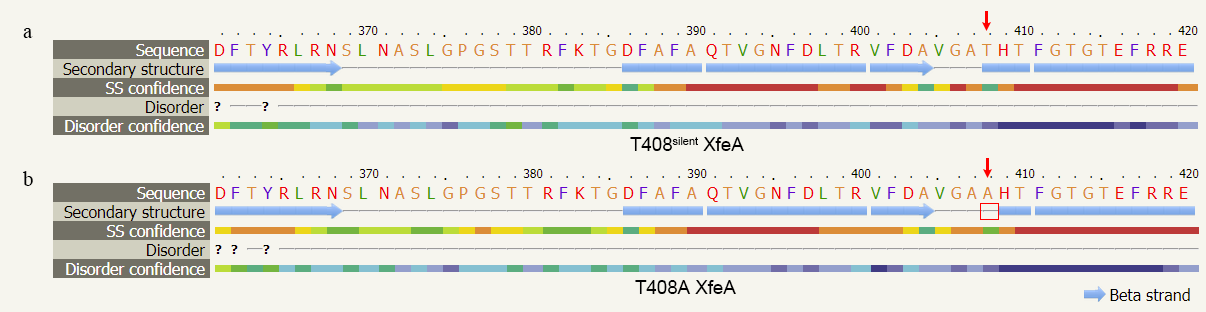

### Fig S6.tif

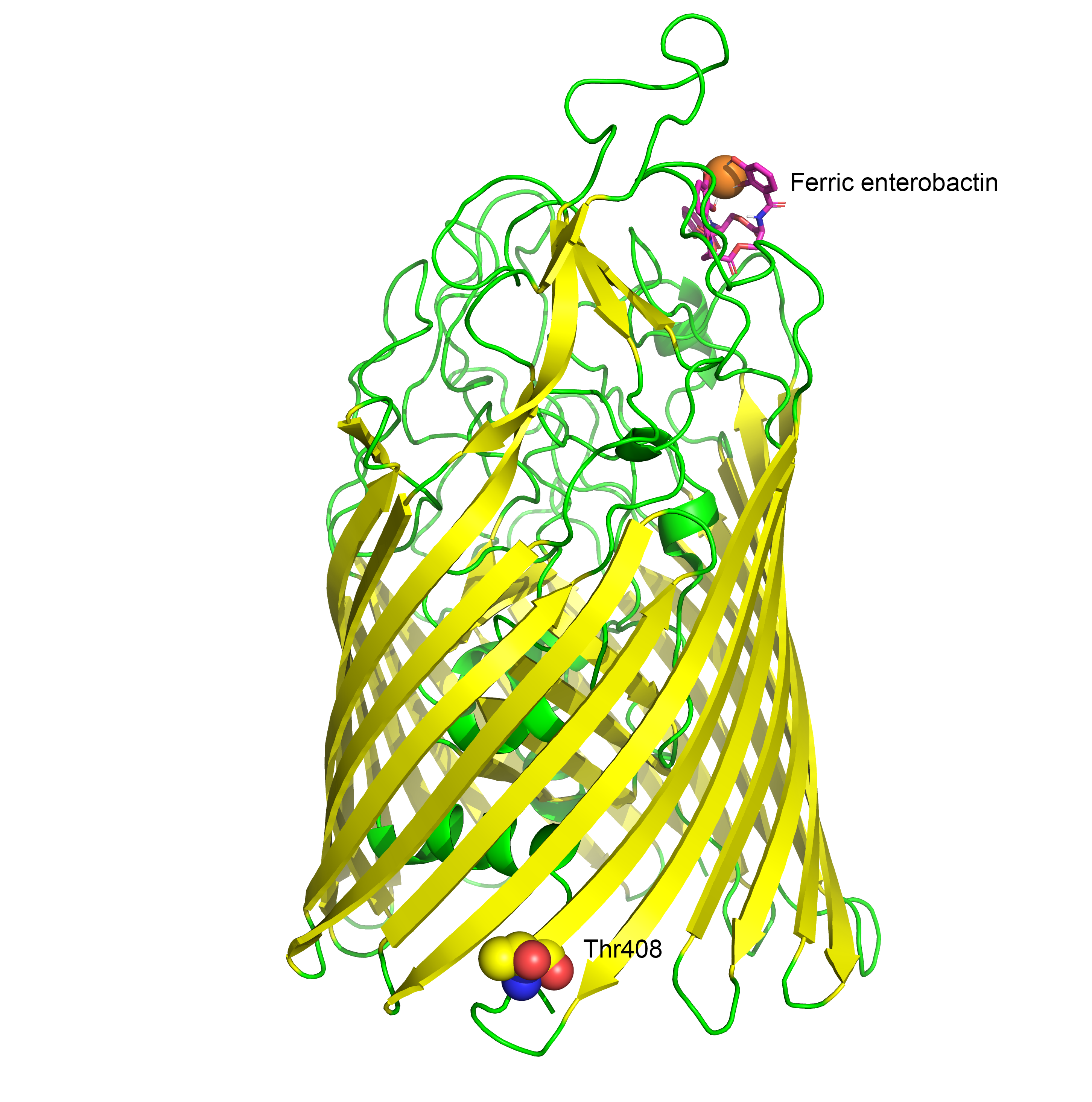
